# Bacterial strain-dependent dissociation of cell recruitment and cell-to-cell spread in early *M. tuberculosis* infection

**DOI:** 10.1101/2022.05.10.491435

**Authors:** B. Shoshana Zha, Ludovic Desvignes, Tawania J. Fergus, Amber Cornelius, Tan-Yun Cheng, D. Branch Moody, Joel D. Ernst

**Author notes:** **Corresponding Author:** Joel Ernst, MD 1001 Potrero Ave, Room 601 UCSF Box 1234. Co-first authors; the order was determined by the leadership role in writing the paper (B. Shoshana Zha, Ludovic Desvignes). Abbreviations: Early secretory antigenic target secretion system (ESX), Mediastinal lymph node (MdLN), Mononuclear phagocyte (MNP), Monocyte-derived dendritic cells (moDC), Monocyte-derived recruited macrophages (RM), *Mycobacterium tuberculosis* (*M. tuberculosis*), neutrophils (neut), Phthiocerol dimycocerosates (PDIM), Phenolic glycolipids (PGL).

## Abstract

In the initial stage of respiratory infection, *Mycobacterium tuberculosis* traverses from alveolar macrophages to phenotypically diverse monocyte-derived phagocytes and neutrophils in the lung parenchyma. Here, we compare the *in vivo* kinetics of early bacterial growth and cell-to-cell spread of two strains of *M. tuberculosis*: a lineage 2 strain, 4334, and the widely studied lineage 4 strain H37Rv. Using flow cytometry, live cell sorting of phenotypic subsets, and quantitation of bacteria in cells of the distinct subsets, we found that 4334 induces less leukocyte influx into the lungs but demonstrates earlier population expansion and cell-to-cell spread. The earlier spread of 4334 to recruited cells, including monocyte-derived dendritic cells, is accompanied by earlier and greater magnitude of CD4^+^ T cell activation. The results provide evidence that strain-specific differences in interactions with lung leukocytes can shape adaptive immune responses *in vivo*.

**IMPORTANCE:** Tuberculosis is a leading infectious disease killer world-wide and is caused by *Mycobacterium tuberculosis*. After exposure to *M. tuberculosis,* outcomes range from apparent elimination to active disease. Early innate immune responses may contribute to differences in outcomes, yet it is not known how bacterial strains alter the early dynamics of innate immune and T cell responses. We infected mice with distinct strains of *M. tuberculosis* and discovered striking differences in innate cellular recruitment, cell-to-cell spread of bacteria in the lungs, and kinetics of initiation of antigen-specific CD4 T cell responses. We also found that *M. tuberculosis* can spread beyond alveolar macrophages even before a large influx of inflammatory cells. These results provide evidence that distinct strains of *M. tuberculosis* can exhibit differential kinetics in cell-to-cell spread which is not directly linked to early recruitment of phagocytes but is subsequently linked to adaptive immune responses.

## INTRODUCTION

*Mycobacterium tuberculosis* is a facultative intracellular bacterium that resides in tissue mononuclear phagocytes (MNP) and granulocytes (1, 2). Previous investigations have shown that *M. tuberculosis* first enters alveolar macrophages (3, 4). After replication in alveolar macrophages, the bacteria subsequently spread to multiple subsets of phagocytes in the lung parenchyma, largely recruited from circulating monocytes (5–7). Subsequently, monocyte-derived lung dendritic cells (moDC) acquire the bacteria and transport them to draining lymph nodes (2), where they transfer antigens to lymph node resident dendritic cells that prime antigen-specific T cells (8, 9). After proliferating and differentiating, effector T cells traffic to the lungs and arrest progression of the infection (10). *M. tuberculosis* antigen-specific T cell responses require 14-17 days to develop after aerosol infection of mice with the commonly-used H37Rv strain (2, 10), and an average of 6 weeks is required for development of adaptive immune responses after infection in humans (11, 12). Considerable evidence indicates that the rate-limiting step in initiating *M. tuberculosis* antigen-specific CD4 T cell responses is acquisition of the bacteria by dendritic cells in the lung, and transport of live bacteria to local draining lymph nodes (8, 10, 13–16).

The mechanisms of cell-to-cell spread of *M. tuberculosis* are incompletely understood, although evidence suggests a role for necrosis of infected cells and release of viable bacteria that are subsequently ingested by other cells (17). Virulent strains of *M. tuberculosis* can inhibit apoptosis (18) through multiple mechanisms including upregulation of the antiapoptotic protein Mcl-1 (19), inhibition of NOX2-induced reactive oxygen species formation (20), upregulation of soluble TNF receptor 2 (21), and inhibition of apoptotic envelope stabilization (22), although induction of apoptosis still occurs and is important for control of *M. tuberculosis in vivo* (23). Conversely, virulent *M. tuberculosis* can promote necrosis through mitochondrial transmembrane potential disruption (24), depletion of host cellular NAD^+^ (25), and inhibition of plasma membrane repair with induction of lipoxin A4 (17, 26), among other mechanisms.

The early secretory antigenic target secretion system (ESX)-1, a type VII secretion system encoded by the RD1 locus of *M. tuberculosis* (27, 28), is implicated in multiple mechanisms for its role in *M. tuberculosis* virulence (29–32). These mechanisms include recruitment of macrophages (33), activation of neutrophils (34), and secretion of immunodominant T cell antigens (ESAT-6 and CFP-10) (35–37). In addition, ESX-1 has been linked to induction of macrophage necrosis (38–40). ESX-1’s role in cellular recruitment and induction of cell death leads to the hypothesis that it plays a key role in *M. tuberculosis* cell-to-cell transfer.

The majority of studies of early innate responses and bacterial cell-to-cell transfer focus on *M. tuberculosis* lineage 4 strains, especially H37Rv or Erdman. However, there are 6 other human-adapted lineages (41) which vary genetically, phenotypically, and degree of induction of inflammatory responses *in vitro* (42). Lineage 2, which includes the Beijing family, is thought to have enhanced pathogenicity (i.e. ability to cause disease) and less protection by BCG vaccination (43, 44). The Beijing family is defined by region of difference (RD) 105, and has 5 sublineages (45). Sublineage 207 has demonstrated increased pathogenicity in guinea pigs, and the strain 4334 within this sublineage was the source of more secondary cases of TB than any other in San Francisco, California (45).

Increasing evidence indicates that distinct *M. tuberculosis* strains exhibit differential interactions with innate immune responses, as reflected by differences in cytokine release (46, 47) and monocyte activation (48). The goal of this work was to characterize the kinetics of *M. tuberculosis* growth and cell-to-cell spread early in infection, to compare the results between two distinct strains of *M. tuberculosis*, and to assess the relationship of cell recruitment and bacterial spread to initiation of CD4 T cell responses. We found that H37Rv and 4334 differ in their recruitment of leukocytes to the lungs, spread from alveolar macrophages to other leukocytes, and the kinetics of antigen-specific CD4 T cell activation. Our work demonstrates strain dependent dissociation of inflammatory cell recruitment and bacterial cell-to-cell spread, and that the timing of cell-to-cell spread impacts the dynamics of antigen-specific CD4 T cell responses.

## RESULTS

### Differential early cell dynamics after aerosol infection with M. tuberculosis strains H37Rv, 4334, and H37RvΔRD1

To test the hypothesis that distinct *M. tuberculosis* strains vary in their initial interactions with the host, we quantitated live bacteria in the lungs of mice infected with H37Rv, 4334, or H37Rv lacking the virulence locus RD1 (H37RvΔRD1) which encodes the ESX-1 type VII secretion system (27, 28) implicated in cell recruitment and inflammation (29, 30, 33, 37). Mice were infected via aerosol with 200-300 colony forming units (CFU) of each bacterial strain and analyzed at frequent time points 1-14 days post-infection, a short time frame that is known to be prior to development of detectable T cell responses. H37Rv exhibited a lag of approximately 72h in which the number of bacteria did not increase in the lungs, similar to that reported in other studies (49, 50). In contrast, strain 4334 grew without a similar lag, and exhibited an increase in the lung bacterial burden that was already detectable by day 3 (Fig. 1A). After day 6, the populations of both strains expanded steadily during the first 2 weeks of infection and were equivalent by day 10 post-infection. In contrast to the two ESX-1-replete strains, H37RvΔRD1 experienced a prolonged lag in expansion (28). The three strains also differed in their induction of increased lung cellularity: H37Rv induced the most marked increases in cellularity; 4334 responses were intermediate; and H37RvΔRD1 induced no detectable changes in the number of lung cells in the initial 14 days of infection (Fig. 1B).

**Figure 1.**
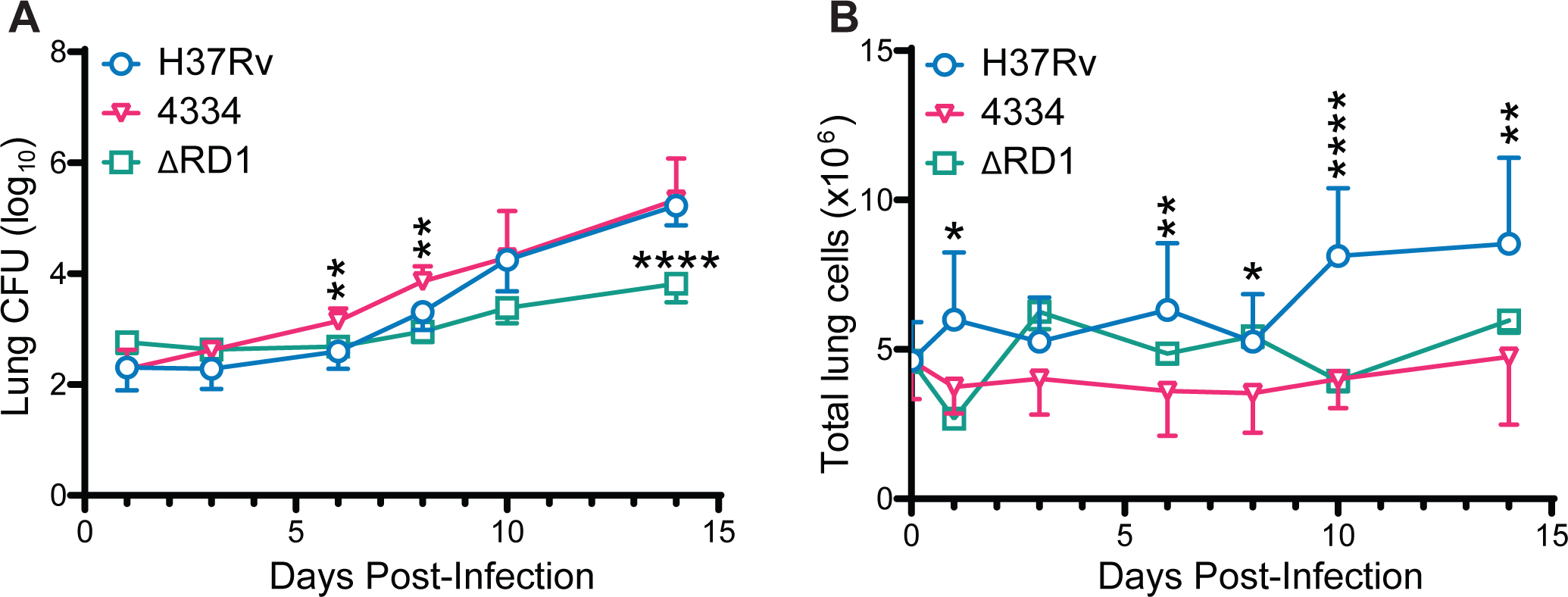
*M. tuberculosis* 4334 grows more rapidly but induces less cell recruitment than H37Rv in early infection. C57BL/6 mice were infected by aerosol with 200-300 Colony Forming Units (CFU) of *M. tuberculosis* H37Rv, H37RvΔRD1 (ΔRD1), or 4334. Lungs were harvested at day 1, 3, 6, 8, 10, and 14 post-infection, and processed to single cell suspensions. (**A**) Total CFU isolated from the lungs of mice infected with the individual strains during the first 14 days of infection. (**B**) Total number of cells isolated from the lungs of mice infected with the individual strains. Results are shown as mean ± SD in 1(ΔRD1), 2 (4334), and 6 (H37Rv) pooled experiments, with n=5 mice per strain per day in each experiment. Statistical significance was assessed by multiple unpaired T-test for each day and Holm-Sidak multiple comparisons correction, with a 95% confidence interval and *p<0.05, **p<0.01, ***p<0.001, ****p<0.0001. For clarity, asterisks above H37Rv and 4334 are comparing only these 2 strains, and below are ΔRD1 compared to H37Rv.

We next compared and characterized the cell populations in the lungs during the first two weeks of infection with the three *M. tuberculosis* strains. For this, we used established cell phenotypic markers and flow cytometry to differentiate subsets in the CD45^+^ cell population: CD11b^neg/lo^CD11c^pos^ alveolar macrophages; CD11b^pos^CD11c^pos^ which includes moDC; Gr-1^hi^CD11c^neg^ neutrophils; Gr-1^int^CD11c^neg^ monocytes; and Gr- 1^neg^CD11c^neg^ monocyte-derived recruited macrophages (RM) (Fig. S1) (6). As shown in Figure 2A, CD11b^neg/lo^CD11c^pos^ alveolar macrophages and CD11b^pos^CD11c^pos^ moDC varied little during the first two weeks and did not differ in lungs of mice infected with 4334 or H37Rv. While Gr-1^hi^CD11c^neg^ neutrophils did not increase in the lungs during the first two weeks of infection with any strain, numbers were consistently lower in the lungs of mice infected with 4334 and H37RvΔRD1 than with H37Rv. The most striking difference between strains was in Gr-1^int^CD11c^neg^ monocytes and Gr-1^neg^CD11c^neg^ RM. As previously reported (2), mice infected with H37Rv showed a progressive increase in Gr-1^int^CD11c^neg^ monocytes and Gr-1^neg^CD11c^neg^ RM in the lungs over the first two weeks, while neither population increased significantly in lungs of mice infected with 4334 or H37RvΔRD1. These findings indicate that the initial cellular inflammatory response to infection with virulent *M. tuberculosis* varies considerably when distinct bacterial strains infect genetically homogeneous hosts.

**Figure 2.**
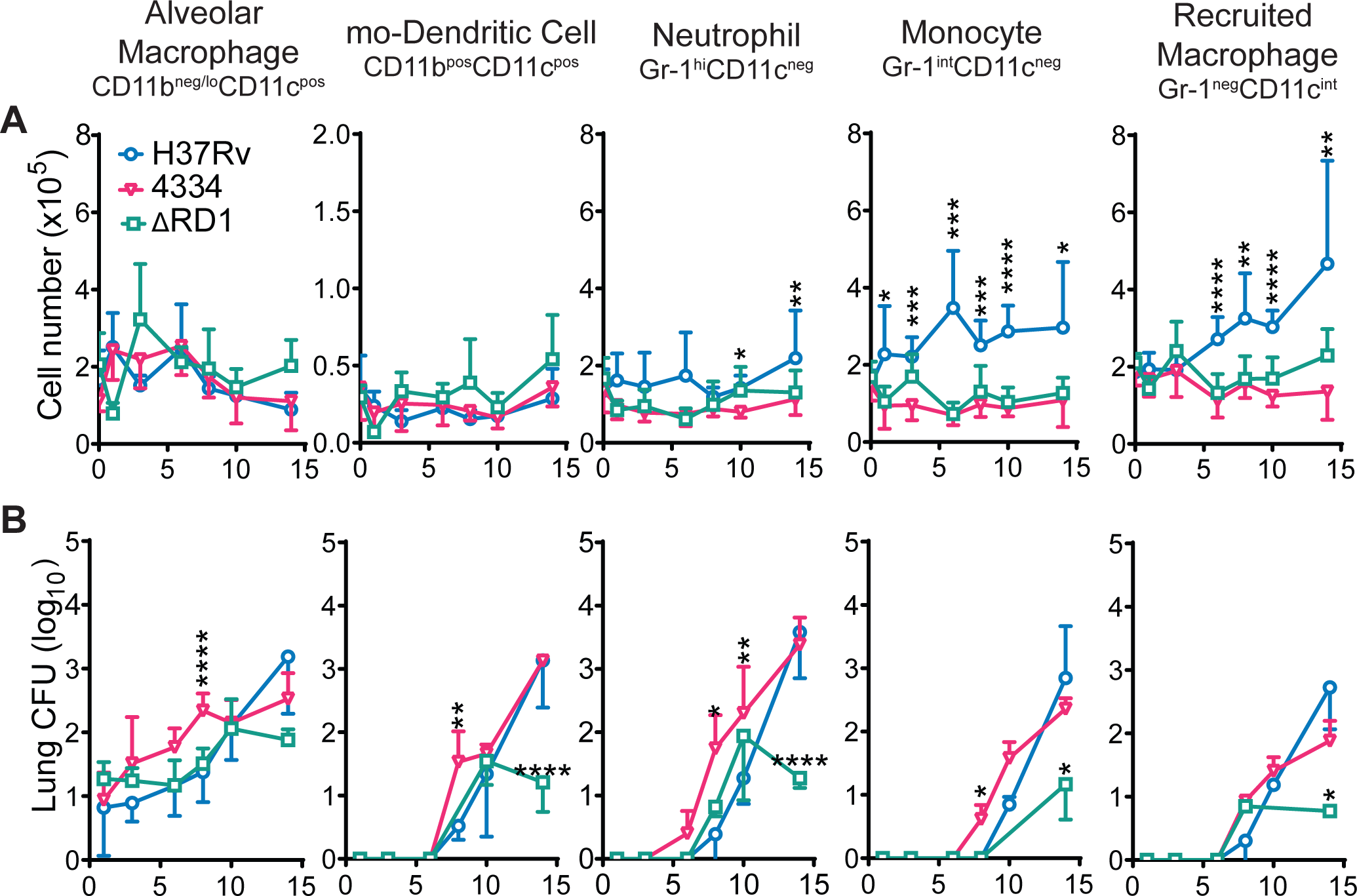
Differential strain-dependent spread of *M. tuberculosis* from CD11b^neg/lo^CD11c^pos^ alveolar macrophages to recruited lung myeloid cells. Mice were infected and lungs harvested as in Figure 1. (**A**) Cells were stained and analyzed by flow cytometry for quantitation of CD11b^neg/lo^CD11c^pos^ alveolar macrophages, CD11b^pos^CD11c^pos^ monocyte-derived dendritic cells, Gr-1^hi^CD11c^neg^ neutrophils, Gr- 1^int^CD11c^neg^ monocytes, and Gr-1^neg^CD11c^neg^ monocyte-derived recruited macrophages in the lungs (see Supplementary Figure 1 and Material and Methods for gating strategy). (**B**) Total CFU in each leukocyte subset after live flow cytometry sorting and plating of sorted cell populations on 7H11 solid media. Results are shown as mean ± SD in 1 (ΔRD1), 2 (4334), and 6 (H37Rv) and pooled experiments, with n=5 mice per strain per day in each experiment. Statistically significant difference between strains was assessed multiple unpaired T-test for each day and Holm-Sidak multiple comparisons correction, with a 95% confidence interval and *p<0.05, **p<0.01, ***p<0.001, ****p<0.0001. For clarity, asterisks above H37Rv and 4334 are comparing only these 2 strains, and below are ΔRD1 compared to H37Rv.

### Mycobacterium tuberculosis strain-dependent dynamics of spread to recruited lung cells

To compare the spread of H37Rv and 4334 from alveolar macrophages to recruited inflammatory cells during the initial phase of infection, we used BSL3- contained flow cytometry sorting of live cells belonging to the previously-identified lung leukocyte subsets (2, 5, 8). We sorted the individual cell populations from infected lungs at frequent intervals and plated the resulting sorted cell fractions on solid media for bacterial quantitation. Consistent with recently-published results generated by flow cytometry detection of cells that harbored fluorescent protein-expressing *M. tuberculosis* (3, 4), we found live bacteria exclusively in alveolar macrophages for the first days post-infection (Fig. 2B, Fig. S2B). On day 6 post-infection, live *M. tuberculosis* were detectable in Gr-1^hi^CD11c^neg^ neutrophils in 2 of the 10 mice infected with strain 4334, but in none of the 15 mice infected with H37Rv, and the average number of live bacteria per cell remained higher in 4334 compared to H37Rv over time (Fig. S2A). Similarly, on day 8 post-infection, 2 of the 10 mice in the group infected with 4334 and none of the 15 infected with H37Rv had detectable live *M. tuberculosis* in Gr-1^int^CD11c^neg^ monocytes, again with higher average number of live bacteria per cell until day 14. Within CD11b^pos^CD11c^pos^ moDC, a significantly higher number of live *M. tuberculosis* were detected for strain 4334 than H37Rv at 8 days post-infection, but the average number of live bacteria per cell in 4334 was higher than H37Rv only until day 10. In Gr- 1^neg^CD11c^neg^ RM, the number of live *M. tuberculosis* per cell was not significantly different between the two strains, although the number of live bacteria per cell was slightly (but not statistically significantly) higher in 4334 than H37Rv at days 10 and 14 post infection.

H37RvΔRD1 expanded in alveolar macrophages at a rate similar to wild-type H37Rv between days 1 and 10. H37RvΔRD1 also spread to CD11b^pos^CD11c^pos^ moDC and Gr-1^hi^CD11c^neg^ neutrophils in quantities comparable to that of wild-type H37Rv by day 10. However, H37RvΔRD1 did not expand further between days 10 and 14. Of note, live H37RvΔRD1 was recoverable in only 2 out of 5 mice in Gr-1^int^CD11c^neg^ monocytes at day 14, and one mouse out of 5 at day 8 in Gr-1^neg^CD11c^neg^ RM.

One known difference with functional importance between *M. tuberculosis* isolates is the presence or absence of phenolic glycolipids (PGL), which are cell wall lipids that modulate virulence (51). PGLs have been implicated in macrophage recruitment (52), dendritic cell uptake, and reduction in inflammatory pathways (53). PGLs are absent from H37Rv and many clinical isolates but are expressed in HN878, a prototypical strain used to study the Beijing family. As 4334 is a member of the Beijing family (44), we determined if 4334 has PGLs. Using electrospray ionization-quadrupole time-of-flight-mass spectrometry (ESI-QTOF-MS), there was no detectable PGL in either 4334 and H37Rv (Fig. 3A,B) under conditions in which PGL was detected in HN878, and the structurally related long chain polyketide phthiocerol dimycocerosates (PDIM) was ionized and detected. This result excludes PGL production as the cause of differential cell-to-cell transfer in these two strains.

**Figure 3.**
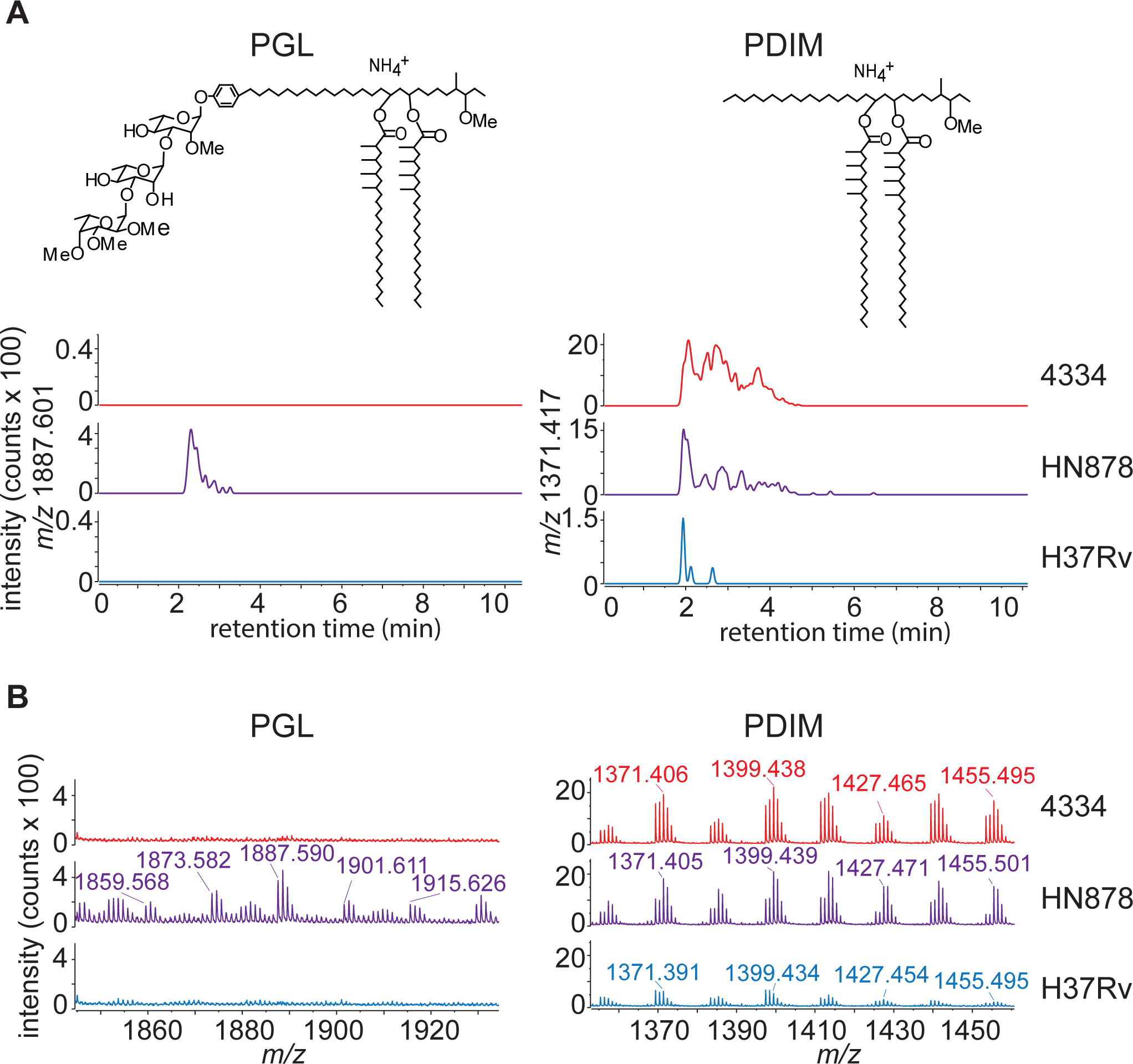
*M. tuberculosis* strains 4334 and H37Rv do not produce PGL. PGL and PDIM analysis of *M. tuberculosis* H37Rv, HN878, and 4334 in 7H9 cultures, using electrospray ionization-quadrupole time-of-flight-mass spectrometry (ESI-QTOF-MS). Shown are representative images of (**A**) extracted ion chromatograms of PGL [C114H212O18+NH4]+ at *m/z* 1887.601 and PDIM [C91H180O5+NH4]+ at *m/z* 1371.417, and (**B**) mass spectra. Strain HN878 was included in the analysis as a known PGL producing positive control.

### *Mycobacterium tuberculosis* 4334 is associated with less alveolar macrophage death than H37Rv

Considering the absence of a lag and immediate expansion of 4334 in alveolar macrophages *in vivo*, we examined the intracellular growth dynamics of 4334 and H37Rv. We infected *ex vivo* murine alveolar macrophages with 4334 or H37Rv at an MOI of 1, and quantitated CFU at 48 and 72h post-infection. In cultured alveolar macrophages, H37Rv did not expand compared with the input by 48h but expanded approximately 5-fold by 72h. In contrast, 4334 expanded 4-fold by 48h, and nearly 10- fold by 72h (Fig. 4A). These growth kinetics mimic the initial phase of infection *in vivo*, in which there was a delay in growth of H37Rv for the first 3 days while 4334 expanded without a similar lag (Fig. 1B).

**Figure 4.**
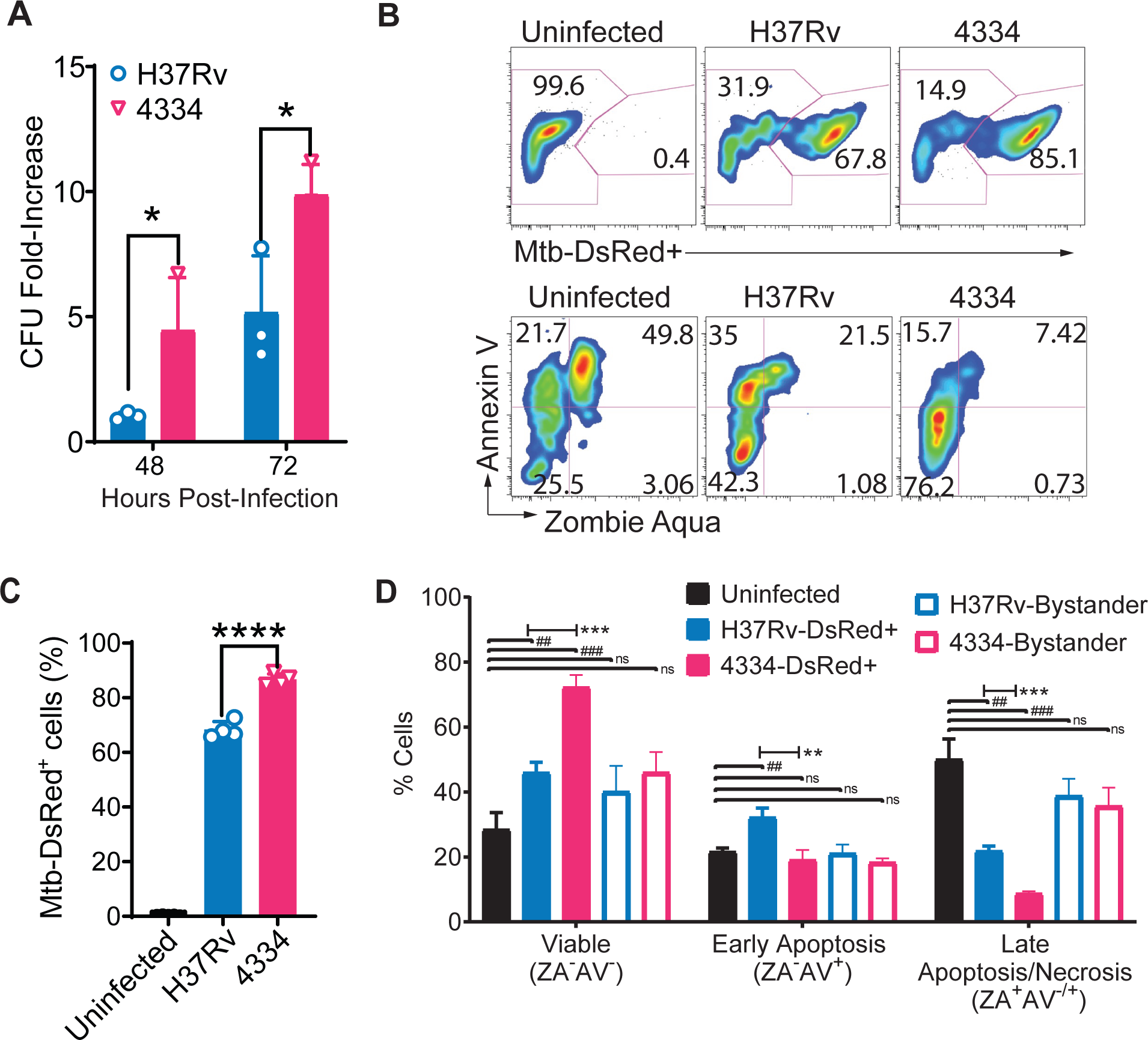
*M. tuberculosis* strain 4334 replicates to a greater extent than H37Rv in alveolar macrophages. (**A**) Cultured alveolar macrophages were infected for 48 or 72h with an MOI of 1 with *M. tuberculosis* H37Rv or 4334. Results are shown as mean CFU fold-increase over initial inoculum. (**B**) Representative flow cytometry analysis of cultured alveolar macrophages infected *in vitro* for 48h with *M. tuberculosis* H37Rv or 4334, each expressing DsRed fluorescent protein and stained with markers of necrosis (Zombie Aqua, ZA) and apoptosis (Annexin V-APC). The upper panels show the frequency of infected cells (DsRed^+^) and lower panels staining with ZA and AV for cells infected with the indicated strain 48h post-infection. (**C**) Percentage of DsRed^+^ cultured alveolar macrophages as quantitated from flow cytometry analysis using the gating shown in panel B. (**D**) Frequency of cultured alveolar macrophages that are viable (ZA^-^ AV^-^), necrotic (ZA^+^AV^-^), early apoptotic (ZA^-^AV^+^) or late cell death (ZA^+^AV^+^) 48h after infection with the indicated strain expressing DsRed fluorescent protein; bystanders are defined as DsRed negative cells. Results are shown as mean ± SD for n=3 per time point and per strain. Data between cells infected with respective strains were analyzed by unpaired Student’s t-test for each day and Holm multiple comparisons correction, with a 95% confidence interval. Asterisks H37Rv vs 4334 (*p<0.05, **p<0.01, ****p<0.0001), hashtag uninfected vs H37Rv or 4334 (^#^p<0.05, ^##^p<0.01, ^###^ p<0.001).

To determine if the apparent difference in growth of 4334 and H37Rv in alveolar macrophages was due to differential cell death, we quantitated infection, early apoptosis, and late cell death 48h post-infection. Using *M. tuberculosis* strains expressing DsRed fluorescent protein and flow cytometry, we observed a higher frequency of infection in cultured alveolar macrophages by 4334 than H37Rv (Fig. 4B,C), consistent with the results obtained by CFU plating. We used a viability marker (Zombie Aqua, ZA) and a phosphatidylserine-binding marker (Annexin V, AV) to quantitate the viability of alveolar macrophages infected *ex vivo* (Fig. 4B). In this scheme, ZA^-^/AV^-^ cells are viable, ZA^-^/AV^+^ cells are considered apoptotic, ZA^+^/AV^-^ are necrotic, and ZA^+^/AV^+^ represent late death, regardless of the mode of death (54). We found significantly less cell death in alveolar macrophages infected by either strain compared to uninfected controls, with quantitatively lower rates of cell death with 4334 compared to H37Rv. Moreover, infection with strain 4334 was associated with a lower frequency of apoptotic cells than H37Rv (Fig. 4D). There was no significant difference in apoptosis or late death between the two strains in bystander cells (that are defined as cells in the same well but lacking DsRed-expressing bacteria).

Cell-to-cell spread of *M. tuberculosis* is thought to involve bacteria released by dying cells. However, our results indicate that 4334 spreads more efficiently despite being associated with lower rates of death of alveolar macrophages. We therefore sought alternative explanations for the early spread of 4334. Since the ESX-1 type VII secretion system has also been implicated in cell-to-cell spread of *M. tuberculosis*, we used liquid chromatography-mass spectrometry (LC-MS/MS) to analyze culture filtrate (i.e. secreted) proteins of 4334 and H37Rv. This revealed significantly higher levels of 3 ESX-1 substrates (EspA, EspB, EspF) and relative higher levels of 3 additional substrates (EspC, ESAT-6, CFP-10) in culture filtrates of 4334 compared with H37Rv (Fig. S3). In contrast, proteins secreted by non-ESX mechanisms (Ag85B and Ag85C) did not differ between the two strains. The finding that cell-to-cell spread of *M. tuberculosis* is associated with quantitative ESX-1 activity and not with the frequency of cell death suggests an alternative mechanism of cell-cell spread and dissemination of *M. tuberculosis* independent of cell death.

### Mycobacterium tuberculosis strain-dependent dynamics of antigen-specific CD4 T cell priming

*M. tuberculosis* induction of antigen-specific CD4 T cell responses requires transport of live bacteria by migratory moDC from the lungs to the mediastinal draining lymph node (8, 10, 13). Since we found more live bacteria in CD11b^pos^CD11c^pos^ moDC from mice infected with 4334 than H37Rv as early as day 8 post-infection (Fig. 2B), we hypothesized that this could result in earlier antigen-specific CD4 T cell priming in the lung-draining lymph node. We first quantitated live bacteria in the mediastinal lymph node (MdLN) on day 14 post-infection, corresponding to the onset of antigen-specific CD4 T cell priming (10). This revealed approximately 3-fold more 4334 than H37Rv in the lymph node (Fig 5A). We have previously established that a threshold number of bacteria are required in the MdLN for priming of *M. tuberculosis* antigen-specific CD4 T cells (10), and therefore hypothesized that 4334 might activate an earlier CD4 T cell response than H37Rv. To test this, we adoptively transferred *M. tuberculosis* Ag85B- specific TCR transgenic (P25TCR-Tg) naïve CD4 T cells (labeled with CellTrace Violet) into mice 24h prior to aerosol infection. Examination of the adoptively transferred T cells isolated from the MdLN revealed no proliferation at day 10 post-infection in mice infected with either strain. In contrast, by day 14 post-infection, P25TCR-Tg CD4 T cells had proliferated and expanded in mediastinal lymph nodes of mice infected with strain 4334 but not H37Rv (Fig. 5B-C). At day 17, P25TCR-Tg CD4 T cells had also proliferated in mice infected with H37Rv, although at significantly lower frequency than in mice infected with 4334.

**Figure 5.**
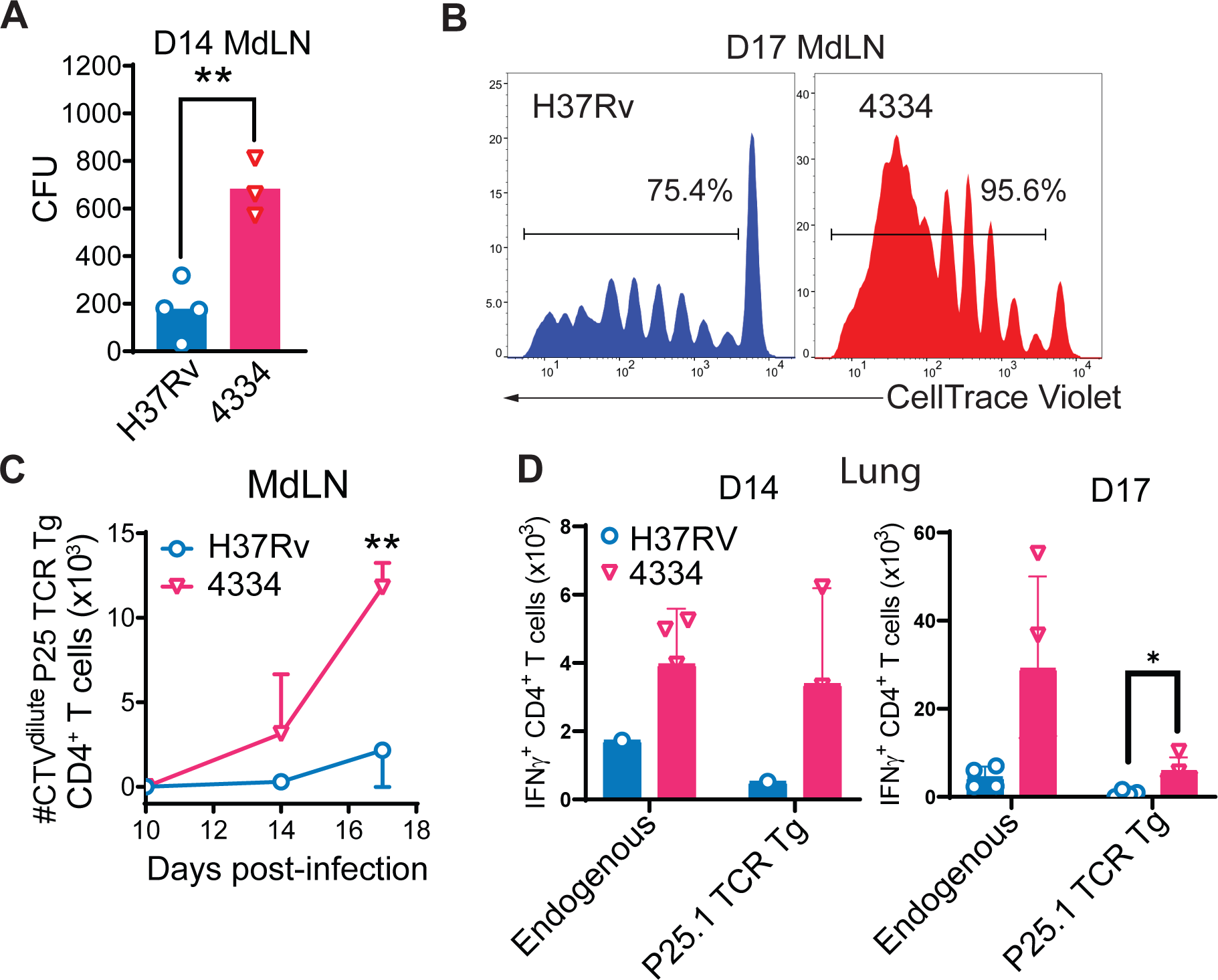
Earlier dissemination of *M. tuberculosis* strain 4334 to lymph nodes is associated with a greater magnitude of Ag85B-specific CD4 T cell priming. (**A**) Bacterial load in mediastinal lymph nodes (MdLN) of mice infected 14 days earlier with *M. tuberculosis* H37Rv or 4334. Results are shown as mean CFU fold-increase over initial inoculum. Flow cytometry assessment (**B**) and absolute quantitation (**C**) of the proliferation of CellTrace Violet-labeled adoptively transferred naïve P25TCR- Tg/CD45.1 CD4^+^ T cells in the MDLN 10-, 14-, and 17-days post-infection with H37Rv or 4334. Percentages shown in panel B are the frequencies of adoptively transferred P25TCR-Tg CD4^+^ T cells in which CTV was diluted as a result of cell division (CTV^dilute^). Results in panel C are shown as mean ± SD, for n=4 mice per day and per strain. (**D**) Quantitation of IFNγ^+^ endogenous and P25TCR-Tg (as defined by congenic markers CD45.1/2) CD4^+^ T cells in the lungs of mice infected for 14 or 17 days with H37Rv or 4334. Results are shown as mean ± SD, for n=4 mice per day and per strain. Flow cytometry analysis for these results is shown in Supplementary Figure 4B. Differences between *M. tuberculosis* strains were assessed by Student’s t-test for each day and Holm multiple comparisons correction, with a 95% confidence interval and *p<0.05, and **p<0.01.

IFNγ is essential for control of *M. tuberculosis* (55, 56). Therefore, we analyzed both total IFNγ concentrations and CD4 T cell specific IFNγ production in the lungs of infected mice of both strains. We found total IFNγ concentrations in lung homogenate supernatants to be significantly higher in mice infected with 4334 than H37Rv at day 14 post-infection (Fig. S4A). We next quantitated IFNγ-producing T cells in the lungs (Fig. S4B) and found that both endogenous and P25TCR-Tg IFNγ-producing T cells were more abundant in mice infected with 4334 than H37Rv at day 14 post-infection, though the difference was not statistically significant. By day 17 post-infection, P25TCR-Tg but not endogenous, IFNγ-producing T cells were significantly more abundant in mice infected with 4334 than H37Rv (Fig. 5D). Together, these results indicate that *M. tuberculosis* strain 4334 spreads to moDC and initiates antigen-specific CD4 T cell responses earlier than does strain H37Rv. This earlier spread to dendritic cells and T cell priming happens despite strain 4334 inducing fewer inflammatory cells to the lung early in infection, and is associated with accelerated priming of antigen-specific CD4 T cells.

We investigated whether the more robust and earlier T cell response correlated into superior long-term control of infection. Indeed, at 7 weeks post-infection, we found a half-log reduction in CFU recovery from the lungs of mice infected with *M. tuberculosis* strain 4334 versus H37Rv (Fig. S5A). Interestingly, while total cell numbers in the lungs of mice infected with *M. tuberculosis* strain 4334 were significantly lower than H37Rv infected mice (Fig. 2B), by 7 weeks this was reversed (Fig. S5B), and the total number of lymphocytes appears to account for the difference in cellularity (Fig. S5D). The increased cellularity also correlates with more granuloma-like lesions in the lungs of mice infected with 4334 (Fig. S5C, S5E), resembling the pathologic findings reported in guinea pigs (44).

## DISCUSSION

The early host responses to pulmonary infection with *M. tuberculosis* are beginning to be clarified (3, 4), although studies to date have not examined the potential impact of bacterial strain diversity. We analyzed early cellular responses in mice infected by aerosol with 4334, a lineage 2 strain (44), and compared this to responses to H37Rv, from lineage 4. Using flow sorting of live lung cells, we compared cell populations, bacterial growth, and bacterial spread from alveolar macrophages to recruited cells in the first 2 weeks of infection. We found that 4334 induces less inflammatory cell recruitment than H37Rv, resembling that of H37Rv lacking the RD1 locus. However, 4334 spreads beyond alveolar macrophages earlier and is present in greater abundance in monocytes, neutrophils, and moDC. This results in earlier trafficking of bacteria to the mediastinal lymph node, and in turn, is accompanied by earlier and greater magnitude activation of Ag85B-specific CD4 T cells.

Recent studies have confirmed that alveolar macrophages are the first cells infected in the lungs by *M. tuberculosis*, a fact that had long been speculated (6) but only recently demonstrated (3, 4). Alveolar macrophages are initially permissive to *M. tuberculosis* replication (4), likely through their inability to mount a large inflammatory response while leading to the upregulation of self-preserving Nrf2 pathways (4) and lipid metabolism (57). We did not find a difference in total CD11b^neg/lo^CD11c^hi^ alveolar macrophages cell numbers after infection. However, there is an earlier expansion of 4334 with significantly higher CFU 8 days post-infection than in H37Rv infected mice, due to a lag in initial H37Rv growth *in vivo* (49, 50). Since alveolar macrophages were the only infected cells during this initial phase, we hypothesized that the differential expansion of the two strains could be due to differences in intracellular replication in alveolar macrophages. We confirmed that 4334 replicates at a higher rate than H37Rv in cultured alveolar macrophages, and this is accompanied by a lower rate of cell death.

By 2 weeks after infection, alveolar macrophages have been found to translocate from the alveolar space to the lung interstitium (3), providing the opportunity for spread of *M. tuberculosis* to other cells. Extending our previous work as well as that of others (3–5, 8), we observed spread of both H37Rv and 4334 to monocyte-derived dendritic cells and macrophages. Here, we found differential spread to the CD11b^pos^CD11c^pos^ population, with 4334 appearing in this population at a higher rate than H37Rv 8 days post-infection. Notably, this cell population includes moDC (5, 6), and although a small percentage of alveolar macrophages also express CD11b (58, 59) with potential upregulation during inflammation and infection (3, 60), those cells have not been found to migrate to lymphoid tissues. Nonetheless, with the combined findings of an earlier and higher rate of spread to neutrophils and monocyte-derived cells in mice infected with 4334 versus H37Rv, our data implies greater spread of 4334 from alveolar macrophages to recruited cells in spite of inducing recruitment of fewer inflammatory cells than H37Rv.

The determinants and mechanisms of *M. tuberculosis* cell-to-cell spread are not well understood, especially *in vivo*, but likely depend on multiple factors including host cell recruitment, intracellular bacterial growth, cellular release, and survival in extracellular spaces. In this work, we found that while 4334, a lineage 2 strain with increased pathogenicity in guinea pigs (44) and the ability to induce higher levels of type I interferons (61), recruits significantly fewer monocytes and cells that differentiated from monocytes but demonstrates superior growth and/or survival rates in the cells it does infect compared to H37Rv. This suggests that progression of infection is not solely determined by cellular recruitment but also by the rates of intracellular replication and transfer between host cells. Differential growth dynamics between *M. tuberculosis* strains has been shown in multiple primary cell types, including human monocytes (62), human monocyte-derived macrophages (63), and murine bone marrow-derived macrophages (64). We hypothesize that the permissiveness of alveolar macrophages for *M. tuberculosis* growth in the initial stage of infection (4, 57) maximizes the intracellular growth variations between strains.

Multiple mechanisms can determine the intracellular survival and growth of *M. tuberculosis*. We focused on differential host cell death since this is thought to play a key role in cell-to-cell spread. Death-receptor induced apoptosis (23) and subsequent efferocytosis (65) are key host mechanisms for control of *M. tuberculosis*. As opposed to avirulent strains lacking the operon RD1 and thus functional ESX-1 machinery, virulent strains of *M. tuberculosis* have been found to inhibit apoptosis (18) and promote necrosis (66) through multiple mechanisms. Necrosis is hypothesized to be the major mode of host cell death for bacterial cell-cell spread, as it allows release of bacilli into the extracellular space for phagocytosis by permissive host cells, rather than being contained in apoptotic vesicles and killed through efferocytosis (17, 33, 67, 68). In contrast to that model, we found that 4334 sustains alveolar macrophage *ex vivo* survival to a greater extent than does H37Rv, yet 4334 spreads more readily to other cells in the lungs.

ESX-1 is implicated in the induction of cell death through the action of secreted proteins including ESAT-6 (40), which is concentration dependent. We have shown that 4334 secretes higher levels of ESX-1 substrates compared to H37Rv, but this is not correlated with activation of cell death of infected alveolar macrophages *ex vivo*. One hypothesis of these discordant findings is the limitation of using *ex vivo* infection modeling, however, the significant protection of alveolar macrophage viability in 4334- infected cells over H37Rv-infected cells argues against differential induction of cell death as a means of cell-to-cell spread. Rather, these results are compatible with a model in which nonlytic release contributes to cell-to-cell spread of *M. tuberculosis* in the lungs (70). Nonlytic release has been demonstrated in *Dictyostelium* amoebae, where *M. marinum* and *M. tuberculosis* egress via ejection rather than lysis of the amoebae (71), and *in vivo* in zebrafish where nonlytic cell-to-cell transfer of *M. marinum* has been observed (72). Furthermore, *M. tuberculosis* can grow extracellularly (73, 74), adding an additional possibility that 4334 survives extracellularly more effectively than H37Rv, allowing uptake by diverse phagocytic cells. Teasing these differenitals is of essential importance to understand the mechanism of tuberculosis cell spread.

We also found significantly more live 4334 *M. tuberculosis* than H37Rv in the draining mediastinal lymph node 14 days post-infection. This correlates with the timing and speed of 4334 spread to moDC, which transport *M. tuberculosis* to the draining mediastinal lymph node (2, 9). Of note, we focused on CD11b^pos^CD11c^pos^ DCs in this study. During homeostasis, conventional DCs (cDCs) have been classified into two populations in mice: cDC-1 (CD26^+^CD11c^+^CD103^+^CD11b^-^) and cDC-2 (CD26^+^CD11b^+^CD11c^+^CD172a^+^) (75, 76). cDC-1 have been characterized as inducing CD8 T cell activation (77) while cDC-2 are implicated in CD4 T cell activation (78). In previous work, we found that cells resembling cDC-2 carry live *M. tuberculosis* to the draining mediastinal lymph node (2, 5), and cDC-1 do not appear to be significantly infected by *M. tuberculosis* in the first two weeks of infection (4). We have previously shown cDC-2-like cells are derived from monocytes (5), and therefore have termed this population moDC. However, CD11b^pos^CD11c^pos^ migratory DCs do not prime CD4 T cells efficiently during *M. tuberculosis* infection (9), and we and others have previously demonstrated that infected moDC transfer antigen to resident mediastinal lymph node DC that in turn activate T cells (8, 9). Here, we observe a significant increase of T cell activation at 17 days post-infection in mice infected with 4334 which correlates with a higher bacterial burden in the draining lymph node at day 14 post-infection. These findings are consistent with the finding of earlier spread of the bacteria to CD11b^pos^CD11c^pos^ moDC in the lungs, but do not indicate whether the greater magnitude of Ag85B-specific CD4 T cell activation in the lymph node is the consequence of more moDC presenting antigen, or more antigen presented per moDC.

Earlier T cell activation can lead to superior control of *M. tuberculosis* infection (13). Here, we demonstrate that 4334 induces an earlier antigen-specific T cell response which is associated with a reduction in CFU 7 weeks post-infection compared to H37Rv infected mice. In addition, the quantitatively greater accumulation of lymphocytes in the lungs of mice infected with 4334 is associated with an increased number of inflammatory lesions in the lung at this time point.

A limitation of this work is the resolution of the flow sorting strategy. As the work prioritized identification and characterization of the cell populations that harbor live *M. tuberculosis* in the initial days after infection when the bacterial burdens are low, the number of surface markers used for sorting and the resolution of certain cell subsets was limited. Nevertheless, the resolution of the subsets was sufficient to allow tracking of the spread of *M. tuberculosis* from alveolar macrophages to recruited inflammatory cells over time. An additional limitation is that the genetic and molecular differences between H37Rv and 4334 were not identified, although we do rule out the lineage 2 specific virulence lipid, PGL, as one potential difference. While both strains were isolated from patients with pulmonary TB, H37Rv was initially isolated in 1905 (79), and 4334 was isolated in the mid-2000s (44). It is not known whether or how passage and storage of the two strains has resulted in genotypic or phenotypic differences from the original patient isolates. However, we do rule out the lineage 2 specific virulence lipid, PGL, and comparative genetic analysis of these strains, as well as future work on the recognized differences between lineage 2 and 4 strains can now be considered in light of these detailed findings during early in vivo infection.

In conclusion, we found that *M. tuberculosis* strain-dependent differences in the rate of spread from alveolar macrophages to recruited leukocytes in the lungs is associated with lower rates of alveolar macrophage death and with a lower rate and extent of initial leukocyte recruitment to the lungs. The finding that enhanced spread from alveolar macrophages is associated with lower rates of alveolar macrophage cell death suggests that nonlytic mechanisms are likely to contribute to cell-to-cell spread of *M. tuberculosis* in vivo. The data also reveal that the strains examined differ in their ESX-1- (type VII) but not SecA (type I)-mediated protein secretion activity, which correlates with enhanced spread, suggesting that Esx-1 may contribute to nonlytic spread of *M. tuberculosis* in the early stages of infection in the lungs. Understanding the mechanisms underlying these findings will provide further insights into the virulence of *M. tuberculosis*.

## METHODS

### Mice and Care

C57BL/6 mice were purchased from the Jackson Laboratory. P25 TCR Tg mice whose CD4 T cells recognize peptide 25 (aa 240-254) of *M. tuberculosis* antigen 85B in complex with mouse MHC II I-A^b^ (10, 80) were crossed in-house with congenic CD45.1^+^ (B6.SJL-Ptprc^a^ Pepc^b^/BoyJ) and with Rag1^-/-^ mice. For infections with *M. tuberculosis*, mice were housed under barrier conditions in the ABSL-3 facility at the NYU School of Medicine. All mice were between 8 and 12 weeks of age at the beginning of the experiment, and mice of both sexes were used. Mice were euthanized by CO_2_ asphyxiation followed by cervical dislocation. All experiments were performed with the prior approval of the NYU Institutional Animal Care and Use Committee (IACUC).

### Cell isolation for *in vitro* and *in vivo* experiments

Alveolar macrophages (AM) were harvested by bronchoalveolar lavage (81), and purity confirmed through flow cytometry analysis with CD11c^+^CD11b^-^ cells (2) above 90% and autofluorescence (82). AM were cultured for up to 4 days in RPMI 1640 with 10% heat-inactivated FBS, 2mM L-glutamine, 1mM sodium pyruvate, 1x β- mercaptoethanol (Gibco), 10 mM HEPES, 10ng/mL of recombinant murine granulocyte-macrophage colony-stimulating factor (Peprotech), and 10 U/ml penicillin – 10 µg/ml streptomycin that was washed out prior to infection.

P25 TCR Tg CD4 T cells were isolated from the secondary lymphoid organs of P25TCR-Tg/CD45.1/Rag1^-/-^ mice by magnetic cell sorting using anti-CD4 conjugated microbeads and an AutoMACS Classic sorter (Miltenyi Biotec), according to the manufacturer’s recommendations.

### *Mycobacterium tuberculosis* strains, growth, and infections

*M. tuberculosis* H37Rv was grown as previously described (83). 4334 was obtained and maintained as previously described (61). Mice were infected via the aerosol route, using an inhalation exposure system (Glas-Col) (2). The infectious dose was quantitated on day 1 by plating whole lung homogenates from 3–5 mice on Middlebrook 7H11 agar. To determine the bacterial load throughout the infection, lungs were harvested, homogenized, and serial dilutions were plated on Middlebrook 7H11 agar. Colony Forming Units (CFUs) were counted after incubation of plates at 37**°**C for 3 weeks.

For *in vitro* infections, mycobacteria were grown to mid-log phase, pelleted at 3750 g, resuspended in PBS + 0.5% Tween 80, re-pelleted and excess Tween 80 washed off by centrifugation in PBS. The final pellet was re-suspended in RPMI 1640 with 10% serum, the bacterial density was evaluated by measuring absorbance at 580 nm, and the multiplicity of infection was adjusted to the mammalian cell density.

### Flow cytometry and cell sorting

Lungs were removed and processed into single-cell suspensions for flow cytometry and cell sorting as previously described (2, 84). Antibodies conjugated to various fluorophores and directed against surface markers were:

| Antibody | Clone | Source |
| --- | --- | --- |
| B220-PE | RA3-6B2 | BioLegend |
| CD4-BUV395 | RM4-5 | BD Biosciences |
| CD4-APC/Cy7 | RM4-5 | BioLegend |
| CD8-APC | 53-6.7 | BioLegend |
| CD8-BV421 | 53-6.7 | BioLegend |
| CD11c-PerCP | HL3 | BioLegend |
| CD11b-APC/Cy7 | M1/70 | BD Biosciences |
| CD19-FITC | 6D5 | BioLegend |
| CD45-PE | 30-F11 | BD Biosciences |
| CD45-FITC | 30-F11 | BD Biosciences |
| CD45.1-AF700 | A20 | BioLegend |
| CD45.2-PerCp/Cy5.5 | 104 | BD Biosciences |
| CD45.2-FITC | 104 | BD Biosciences |
| CD90.2-PerCP/Cy5.5 | 30-H12 | BioLegend |
| Gr-1-APC | RB6-8C5 | BD Biosciences |
| Ly6C-FITC | AL-21 | BD Biosciences |
| Ly6G-AF647 | 1A8 | BioLegend |
| Siglec-F-PE | E50-2440 | BD Biosciences |

For flow cytometry analysis, stained samples were fixed overnight in 1% paraformaldehyde. A minimum of 200,000 events per sample, gated on single cells using forward and side scatter parameters, were acquired using an LSRII and FACSDiva software (BD Biosciences).

For live cell sorting, fluorescently labeled live cells were acquired using a BSL3- contained iCyt/Sony Synergy cell sorter. Cells were first separated in non-hematopoietic cells (CD45^-^), AM (CD45^+^CD11b^-^CD11c^+^), monocyte-derived dendritic cells (moDC, CD45^+^CD11b^+^CD11c^+^) and other myeloid cells (CD45^+^CD11b^+^CD11c^neg/low^). Then, this latter fraction was re-acquired and sorted into neutrophils (Neut, Gr1^hi^CD11c^-^), monocytes (Gr1^int^CD11c^-^) and recruited macrophages (Gr-1^-^CD11c^low^). All cell fractions were collected and plated for CFU recovery and quantitation on agar as described above. For flow cytometry and cell sorting, acquisition data were analyzed using FlowJo software (TreeStar).

### *Mycobacterium tuberculosis*-antigen specific T cell adoptive transfer, proliferation, and IFNγ stimulation

P25TCR-Tg/CD45.1/Rag1^-/-^ CD4^+^ T cells were stained with CellTrace Violet (ThermoFisher Scientific), according to the manufacturer’s recommendations. Cells were re-suspended at 2-3x10^6^ cells/100 µl in sterile PBS for intravenous injection via the retro-orbital route into anesthetized CD45.2^+^ C57BL/6 recipient mice. Recipient mice were infected with aerosolized *M. tuberculosis* 24h post-T cell transfer. T cell proliferation was assessed at selected time points post-infection by quantitating CellTrace Violet dilution by flow cytometry. Whole lung supernatants were analyzed for IFNγ levels by ELISA.

### Quantification of Necrotic and Apoptotic Cells

AM were infected with *M. tuberculosis* expressing DsRed for designated times, after which cells were washed and then lifted with cold PBS containing 0.5 mM EDTA. Cells were fixed, washed, and stained with Stain Zombie Aqua (Biolegend) to quantify necrosis and AnnexinV-APC to quantify apoptosis (BD Biosciences) per manufacturer protocols, washed with PBS, and analyzed by flow cytometry for DsRed (bacteria), Annexin V, and Zombie Aqua.

### Analysis of PDIM and PGL

Bacterial strains were grown shaking at 37°C for ∼7 days to an OD580 of 0.6 in Middlebrook 7H9 broth supplemented with 10% Oleic acid/dextrose/catalase and 0.05% Tween 80. Cultures were centrifuged, supernatant removed, and pellets washed twice, resuspended in 1mL of CH_3_OH and contacted with 25mL CHCl_3_/CH_3_OH (2:1) overnight. Lipid was then extracted, dissolved in CHCI_3_:CH_3_OH at 1mg/mL, and ran on an Agilent Technologies 6520 Accurate-Mass Q-Tof and 1200 series HPLC as previously described (85).

### ESX-1 Secretion Assay

Bacterial strains were grown shaking at 37°C for ∼7 days in Middlebrook 7H9 broth supplemented with 10% Oleic acid/dextrose/catalase and 0.05% Tween 80 to an OD580 0.4 – 0.7. Cultures were centrifuged, washed thrice in Sauton’s media (0.5 g KH2PO4, 0.5 g MgSO4, 4.0 g L-asparagine, 60 ml glycerol, 0.05 g Ferric ammonium citrate, 2.0 g citric acid, 0.1 ml 1% ZnSO4, dH20 to 900 ml, 2.5 ml 20% Tween-80), and incubated for an additional 24 hours. Bacteria were centrifuged at 2500 g for 10 min, and supernatants passed through 0.2 μm syringe filters to obtain filtrates. Samples were reconstituted in 200uL of 2M urea, reduced with DTT at 57°C for 1h, alkylated with 0.5M iodacetamide for 45 min at RT, followed by trypsin digestion and cleansed as previously described (86). After SpeedVac concentration, samples reconstituted in 0.5% acetic acid, and loaded onto an Acclaim PepMap trap column in line with an EASY Spray 50cm x 75µm ID PepMap C18 analytical HPLC column with 2μm bead size using the auto sampler of an EASY-nLC 1000 HPLC (ThermoFisher) and solvent A (2% acetonitrile, 0.5% acetic acid). The peptides were gradient eluted into a Q Exactive (Thermo Scientific) mass spectrometer using a 2h linear gradient from 2% to 40% solvent B (95% acetonitrile, 0.5% acetic acid), followed by 10 min from 40% to 100% solvent B. Solvent B was held at 100% for another 10 min for column wash. Spectra were acquired using the following parameters: resolution of 70,000, an automatic gain control of 1e6, with a maximum ion time of 120 ms, and scan range of 300 to 1500 m/z. Following each full MS scan twenty data-dependent high resolution HCD MS/MS spectra were acquired. All MS/MS spectra were collected using the following instrument parameters: resolution of 17,500, AGC target of 2e5, maximum ion time of 250 ms, one microscan, 2 m/z isolation window, fixed first mass of 150 m/z, and normalized collision energy of 27.

All acquired MS/MS spectra were searched against a combined database for H37Rv and NITR203 on UniProt, using Andromeda search algorithm and MaxQuant for quantitation (87). The data set was filtered to remove proteins with only one unique peptide and that were not detected in all three replicates of at least one strain. LFQ intensity values were log_2_ transformed. A two-sided t-test and correcting for multiple testing by controlling for FDR at 5% using Benjamini-Hochberg’s method. In addition, z-scores were calculated and used to perform hierarchical clustering.

### Histopathology

The left lung was excised, fixed in 10% buffered formalin for one week at room temperature, then embedded in paraffin. 5 μm sections were cut and stained with hematoxylin and eosin. Whole lung sections were scanned at 40X by the NYU School of Medicine Experimental Pathology Research Laboratory, using a Leica SCN400 F whole-slide scanner. Digital images were used for the quantitation of lung histopathology using the open-source image processing software Fiji, with two independent approaches. Briefly, the total surface area of the lung section was calculated, then the contributions of airways and blood vessels to that area were subtracted, leaving the contribution of the structural tissues. The proportion of inflammatory infiltrates within those tissues was then quantitated, either by manual contouring or using automated color density contouring. For each 40X section, the final percentage of lung inflammatory infiltrate was calculated as the average of the values obtained by each method.

### Statistical analysis

Experiments were performed at least twice, with exception of experiments focusing on H37RvΔRD1 and 7 weeks post-infection which were performed once. Results are expressed as mean and standard deviation (SD). Unless otherwise stated, parametric Student two-tailed t test and Holm multiple comparisons correction, with a 95% confidence interval was used to compare experimental groups, with p<0.05 considered significant.

## ACKNOWLEDGMENTS

We thank Diane Ordway and Midori Kato-Maeda for the *M. tuberculosis* strain 4334. Cynthia Portal-Celhay and Thais Klevorn aided in experiments. We acknowledge the expertise and support of Beatrix Ueberheide and Jessica Chapman-Lim at the NYU Proteomics Laboratory, supported in part by NYU Langone Health and the Laura and Isaac Perlmutter Cancer Center Support Grant P30CA016087 from the National Cancer Institute, for proteomics assays. Cell sorting and flow cytometry were performed by Michael Gregory from the NYU Cytometry and Cell Sorting Laboratory, which is supported in part by grant P30CA016087 from the National Institutes of Health/National Cancer Institute. This project was supported by NIH grants R01 AI051242 and R01 AI049313, as well as an award from the Stony Wold-Herbert Fund. L.D. and J.D.E. designed the experiments, B.S.Z, L.D., and J.D.E. analyzed the results; T.J.F., L.D., and A.C. conducted experiments; T.Y and D.B.M. provided the lipidomic analysis and reviewed the manuscript; B.S.Z, L.D., and J.D.E. wrote the manuscript.

## Supplemental Material Legends

**Supplementary Figure 1. Flow cytometry gating strategy for identification of lung myeloid cell populations**. After processing lung tissue into single-cell suspensions and staining, cells were flow sorted first as CD45^neg^ (non-hematopoietic) or CD45^pos^. Further cellular identification occurred using CD11b and CD11c to distinguish between **(1)** CD11b^neg/lo^CD11c^pos^ alveolar macrophages and **(2)** CD11b^pos^CD11c^pos^ monocyte- derived dendritic cells. Further differentiation of CD11b^pos^ cells was done using CD11c and Gr1, as **(3)** Gr1^hi^CD11c^neg^ neutrophils, **(4)** Gr1^int^CD11c^neg^ monocytes, and **(5)** Gr- 1^neg^CD11c^neg^ monocyte-derived recruited macrophages.

**Supplementary Figure 2. Strain-dependent spread of *M. tuberculosis* in recruited myeloid cells.** (**A**) Cells were stained and analyzed by flow cytometry for quantitation of subset and plated on 7H11 media (Figure 2). Depicted are the ratio of the average CFU over the average cell number at each time point obtained post-infection in mice infected with H37Rv, 4344, or ΔRD1. (**B**) Relative frequency of CFU recovered from myeloid cell populations in the lungs of mice infected with *M. tuberculosis* H37Rv, ΔRD1, or 4334 after live flow sorting and plating of sorted cell fractions on 7H11 solid media. Results are shown as mean ± SD in 1(ΔRD1), 2 (4334), and 6 (H37Rv) pooled experiments, with n=5 mice per day and per strain in each experiment.

**Supplementary Figure 3. M. *tuberculosis* strains exhibit differential ESX-1 activity.** H37Rv and 4334 were grown to logarithmic phase in 7H9, resuspended in Sauton’s media for 1 day, and filtrate proteins were reduced, alkylated, and trypsin digested prior to run on LC-MS/MS. Spectra were searched against a combined database for H37Rv and NITR203 on UniProt, using Andromeda search algorithm and MaxQuant for quantitation. Statistical analysis done by Student t-test with Holm multiple comparisons correction, 95% confidence interval, *p<0.05, and **p<0.01.

**Supplementary Figure 4. *M. tuberculosis* strain 4334 induces higher levels of IFNγ in the lungs of mice than H37Rv after two weeks of infection.** (**A**) Total IFNγ levels in the lung homogenate supernatants of mice infected with *M. tuberculosis* H37Rv or 4334. Results are shown as mean ± SD 4 experiments. Statistical significance was assessed by unpaired Student’s t-test for each day and Holm multiple comparisons correction, with a 95% confidence interval and **p<0.01 and ***p<0.001. (**B**) Flow cytometry gating strategy for the quantitation of IFNγ-producing CD4^+^ T cells in the lungs of mice infected with *M. tuberculosis* H37Rv or 4334 for 17 days.

**Supplementary Figure 5. *M. tuberculosis* strain 4334 induces a quantitatively greater adaptive immune response compared to H37Rv that is sustained 7 weeks after initial infection and results in more lesions in the lung.** Mice were aerosol infected with 200-300 Colony Forming Units (CFU) of *M. tuberculosis* H37Rv or 4334 and harvested 7 weeks later. Lungs were processed to single cell suspensions and plated on solid media for total CFU determination **(A)**, counted for total cell number **(B),** and stained for flow cytometry analysis of cellular subsets **(D)** (see Supplementary Figure 1 and Material and Methods for gating strategy). In addition, the left lung of infected mice was fixed and stained with hematoxylin and eosin (H&E). **(C)** Using Fiji open-source software, percentage of lung inflammatory infiltrate was calculated by taking the total surface area of lung, subtracting contributions of airways and blood vessels. Inflammatory infiltrates were quantitated by both manual contouring and automated color density contouring, with the average of these values utilized for the final percentage calculation. **(E)** Representative images of 5 µM sections at 40x (Leica SCN400 F whole-slide scanner) are shown. Results are shown as mean ± SD of 4 mice. Statistical significance was assessed by unpaired Student’s t-test and Holm multiple comparisons correction with a 95% confidence interval and *p<0.05, ***p<0.001, and ****p<0.0001.

## Notes

### Competing Interest Statement

The authors have declared no competing interest.

### Summary of Updates

Supplemental files added

