## Supplementary figures and images for "Bacterial strain-dependent dissociation of cell recruitment and cell-to-cell spread in early *M. tuberculosis* infection"

### Supplemental Figure 1

##### Supplement Figure 1

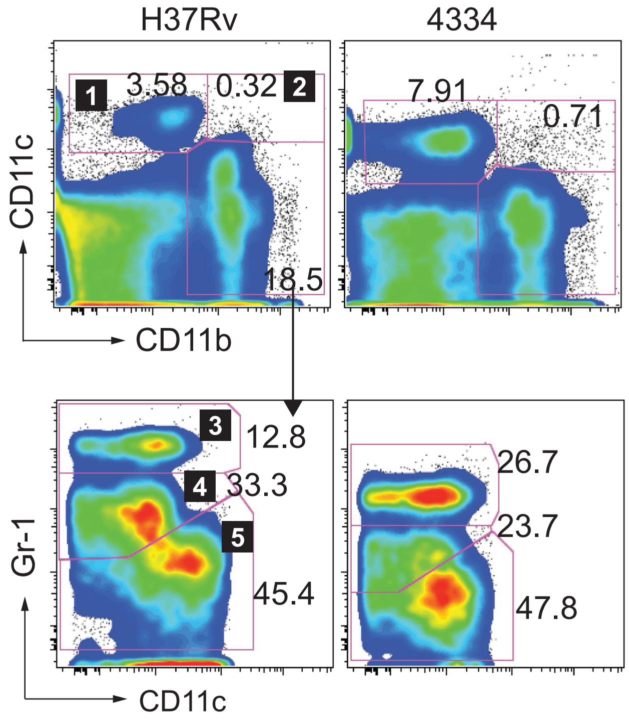

### Supplemental Figure 2

##### Supplement Figure 2

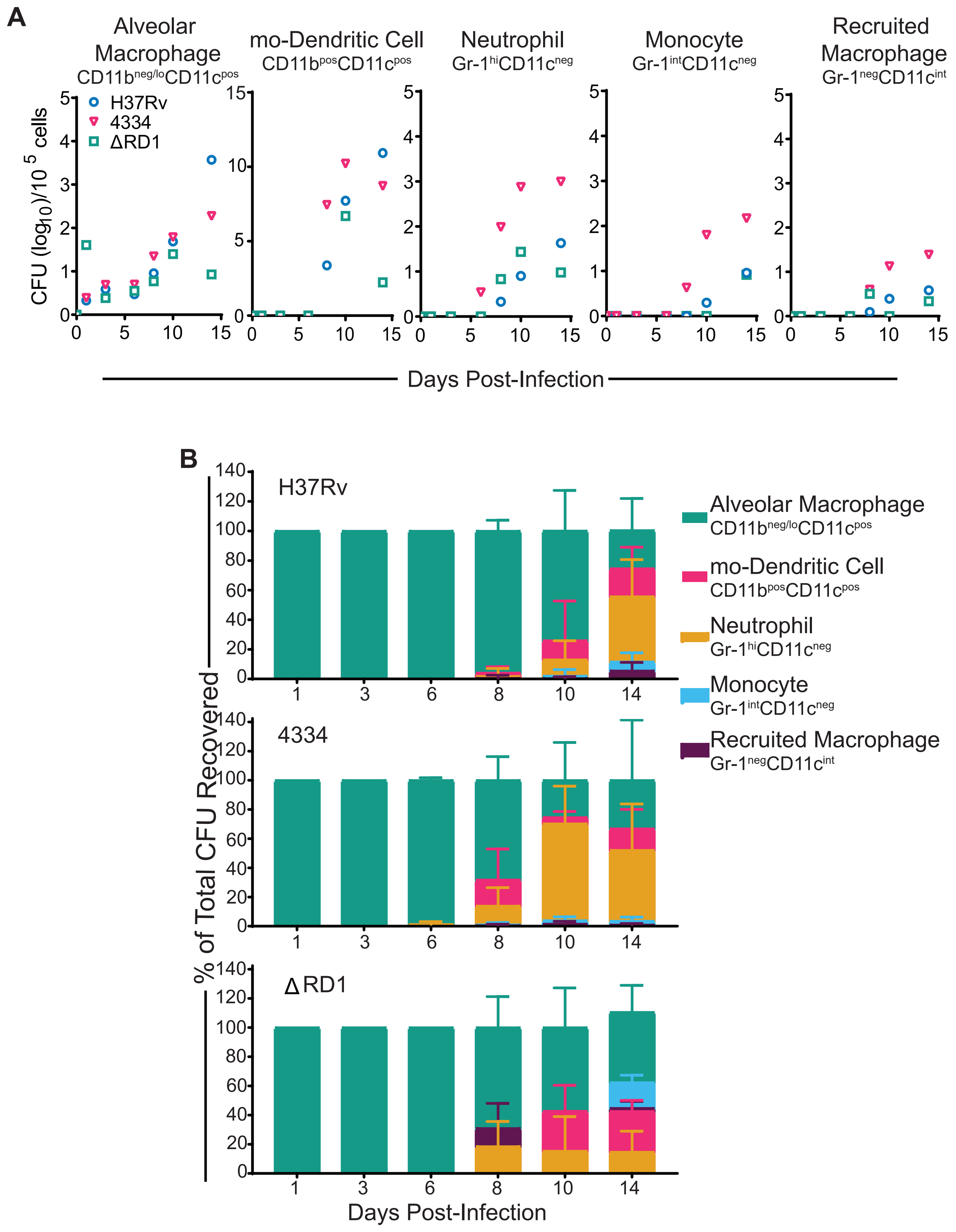

### Supplemental Figure 3

##### Supplement Figure 3

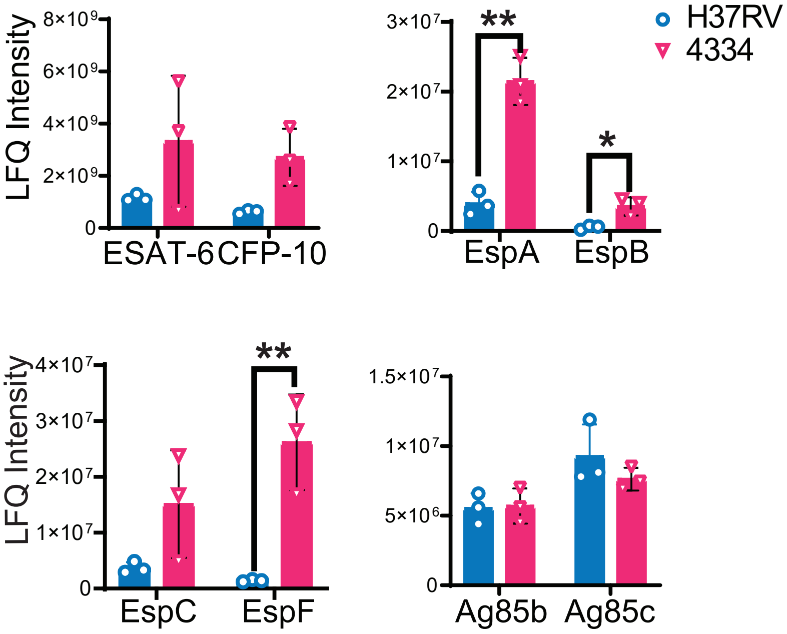

### Supplemental Figure 4

#### Supplement Figure 4

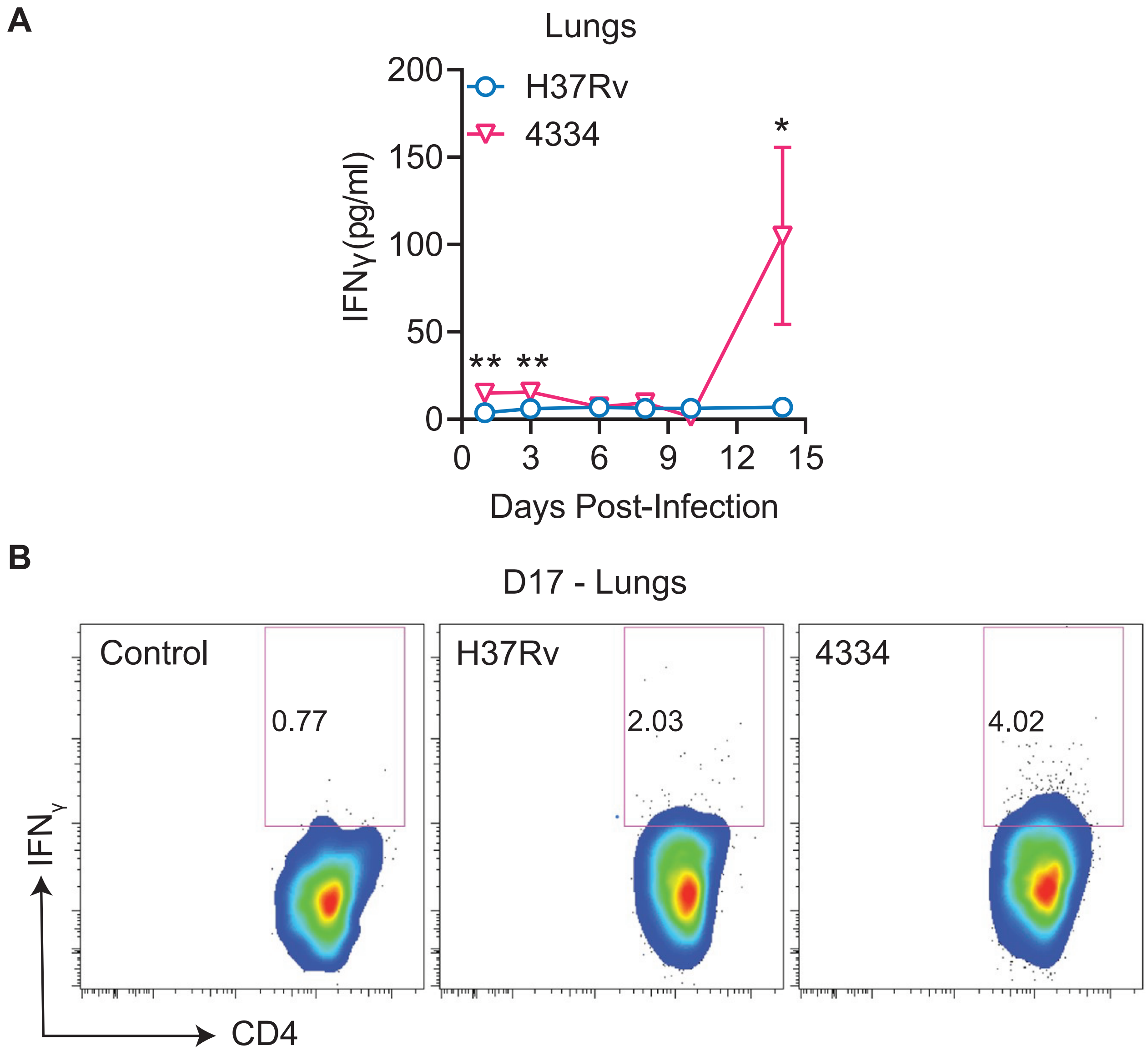

### Supplemental Figure 5

#### Supplement Figure 5

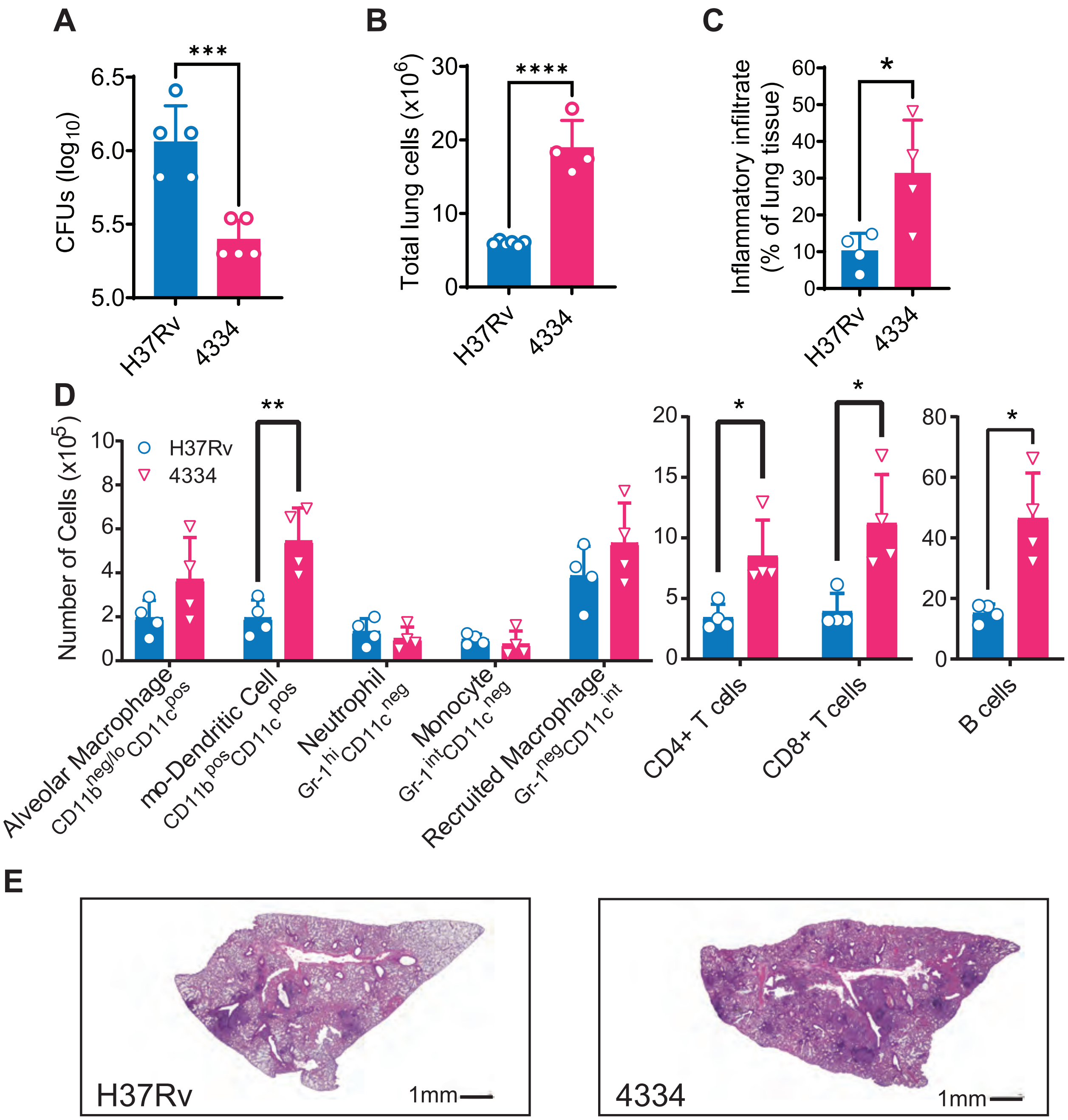
